## Supplemental information for "Piezo1 mechanosensing regulates integrin-dependent chemotactic migration in human T cells"

### for

**by Liu CSC et.al.**

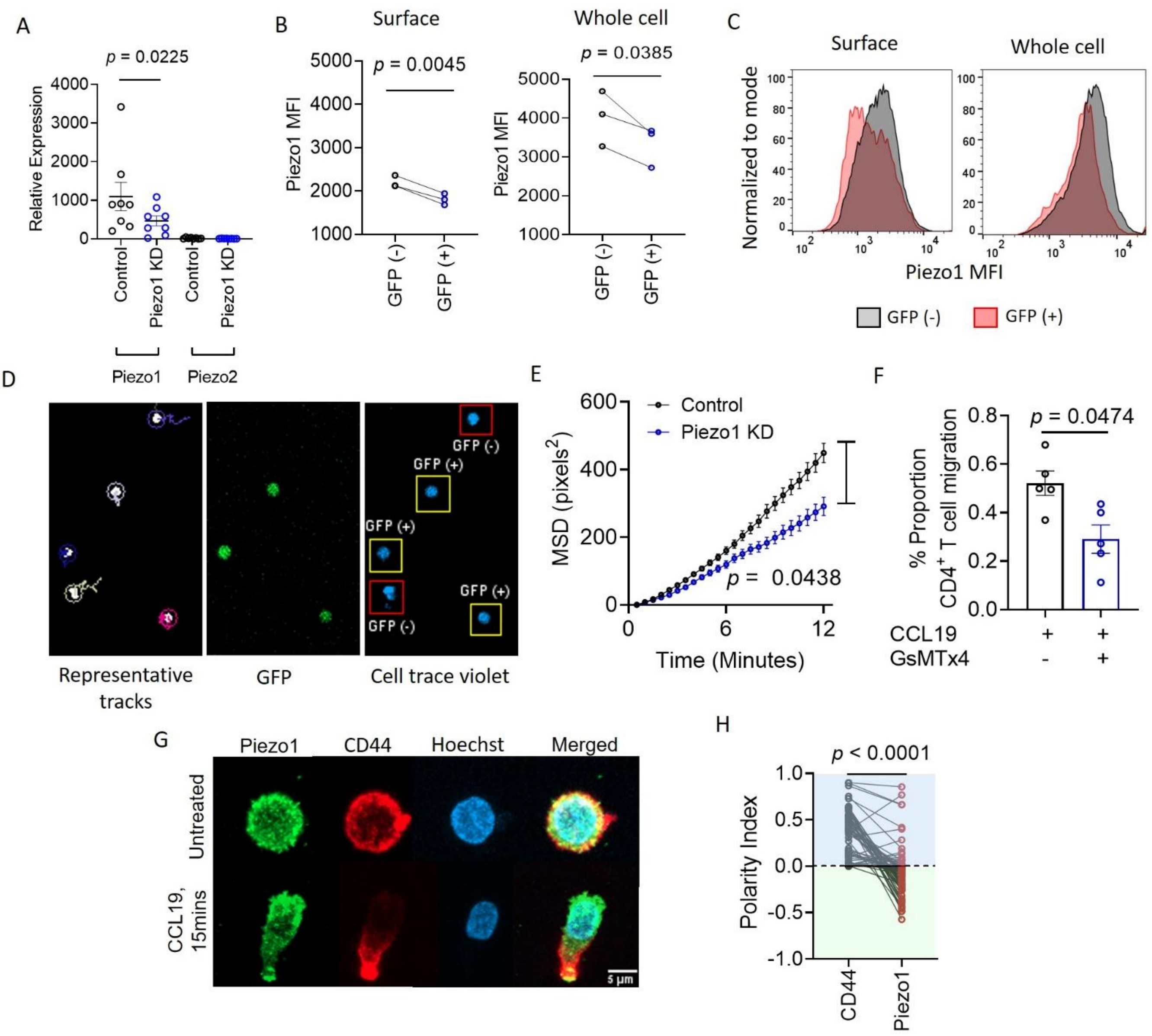


**Figure 1-figure supplement 1**. **A.** Quantitative PCR analysis of Piezo1 and Piezo2 relative expression, in control and Piezo1 siRNA-transfected CD4$^+$ T lymphocytes. **B.** Flow cytometric measurement of Piezo1 mean fluorescence intensity, MFI (left: surface staining; right: intracellular staining) in Piezo1 siRNA and GFP co-transfected cells. **C.** Representative histograms (right two panels) of Piezo1 MFI in GFP (-) and GFP (+) cells in surface stained (left) and intracellularly stained (right) cells. **D.** Representative immunofluorescence image 2D tracks (left), GFP signal (middle) and cell trace violet signal of all cells (right). 20X magnification. E. MSD tracks of control siRNA (n = 357 cells) and Piezo1 siRNA-transfected CD4$^+$ T cells (n = 348 cells) on ICAM1-coated wells in the presence of CCL19. **F.** Effect of Piezo1 inhibition by GsMTx4 on 3D ICAM1-coated transwell migration. **G**. Representative confocal images depicting anti-polarity of Piezo1 and CD44 (uropod) in untreated and CCL19-stimulated conditions. **H**. Polarity indices calculated for Piezo1 and CD44, where negative polarity for Piezo1 means anti-polarity (described in detailed in Methods section). N = 69 cells, each. All data shown is representative of at least 3 independent experiments

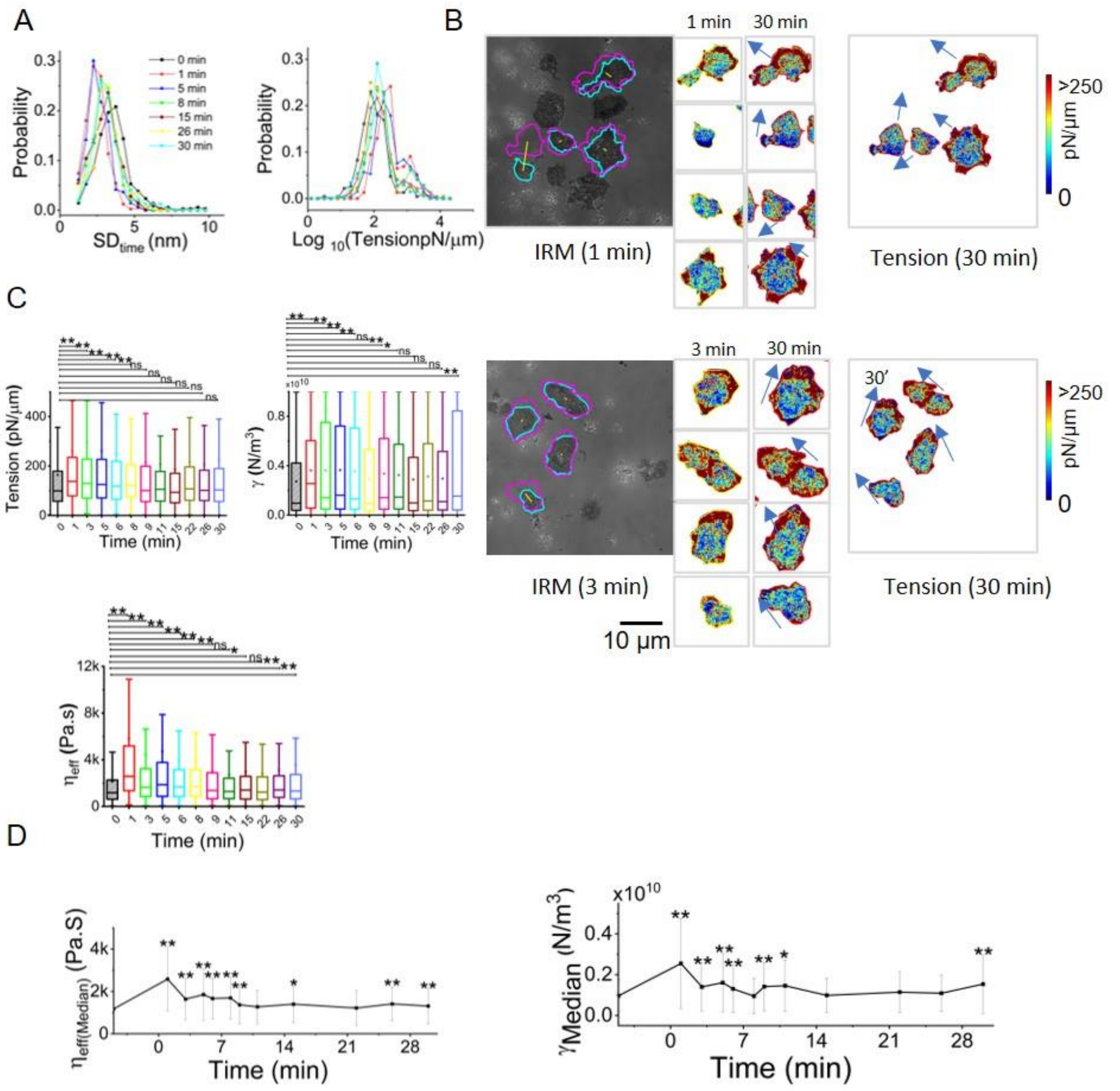


**Figure 2-figure supplement 1. A.** Probability distribution of temporal fluctuation (left) and tension (right) followed over time, **B.** Tension maps showing high-tension edges aligning with direction of cell migration (arrows) before and after addition of chemokine. Cells from 2 fields (top and bottom panel) are depicted in the figure. **C.** Box plots of time-kinetics of tension (upper left), confinement (upper right), effective cytoplasmic viscosity (lower left)**. D.** Temporal trajectory of effective cytoplasmic viscosity (left) and confinement (right) of cells before and after addition of chemokine with ** indicating significant difference from control. ns = *p*>0.0042, **p*<0.0042, ***p*<0.000083 derived using Mann-Whitney U Test with Bonferroni correction. $N_{cells}$ =21, $N_{FBR\ (control)}$ = 1594, $N_{FBR\ (1\ min)}$ =1228, $N_{FBR\ (3\ min)}$ =643, $N_{FBR\ (5\ min)}$ = 1349, $N_{FBR\ (6\ min)}$ =594, $N_{FBR\ (8\ min)}$ =1056, $N_{FBR\ (9\ min)}$ =1562, $N_{FBR\ (11\ min)}$ =638, $N_{FBR\ (15\ min)}$ =1843, $N_{FBR\ (22\ min)}$ =741, $N_{FBR\ (26\ min)}$ =2019, $N_{FBR\ (30\ min)}$ =2746. Scale bar = 10 µm.

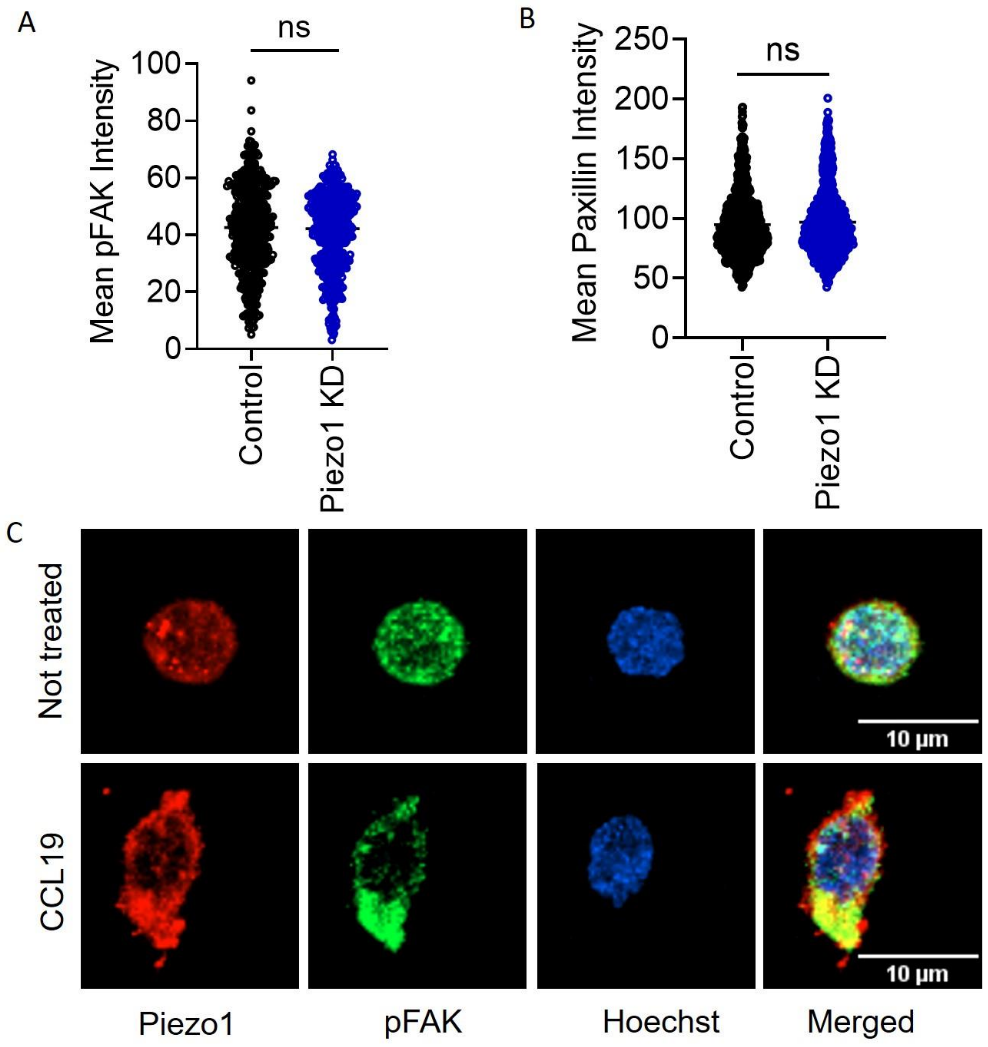


**Figure 3-figure supplement 1. A.** Mean signal intensity of pFAK in CCL19-treated control and Piezo1 KD cells (n > 440 random cells, each). **B.** Mean signal intensity of paxillin in CCL19-treated control and Piezo1 KD cells (n > 780 random cells, each). **C.** Representative images of pFAK and Piezo1 colocalization in untreated (top panel) and CCL19-treated (bottom panel) CD4$^{+}$ T cells.

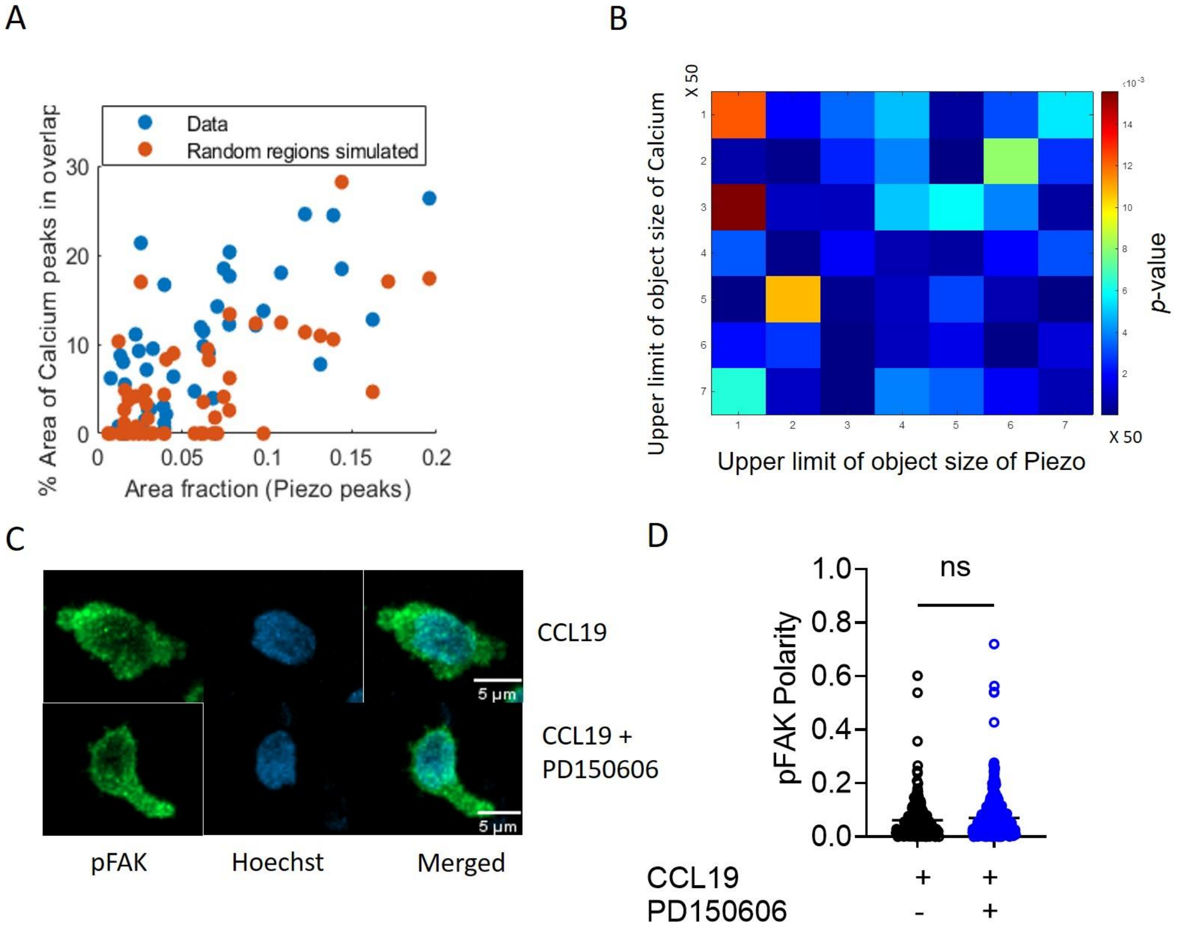


**Figure 5- figure supplement 1. A.** Scatter plot of how percentage area of $Ca^{2+}$ peaks (real and simulated) that overlap with Piezo1 varies with area fraction of Piezo1 showing comparison between real and simulated data. **B.** Color matrix of p-value of comparison of % overlap obtained using real data and simulated data. Color denotes p-value. As 'x' increases maximum object size used for Piezo1 signal changes. As 'y' increases maximum object size used for $Ca^{2+}$ signal changes. **C.** Representative confocal images of human $CD4^{+}$ T cells fixed and stained for phosphorylated FAK. Cells were treated with CCL19 with or without prior incubation with calpain inhibitor-PD150606. **D.** Comparison of pFAK polarity of cells stimulated with CCL19, with or without prior calpain inhibition. n > 230 random cells, each.

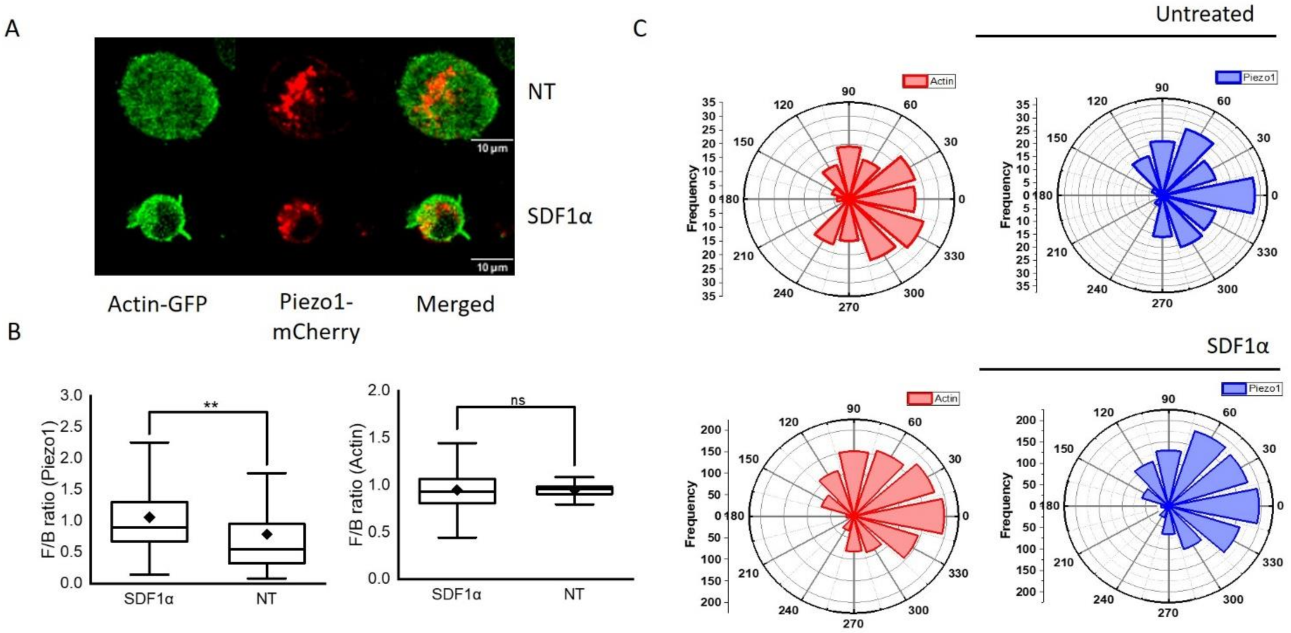


**Figure 6-figure supplement 1. A.** Representative 2D confocal image of Jurkat cell expressing Piezo1 mCherry and actin GFP in the absence (top panel) or presence (bottom panel) of recombinant SDF1α. **B.** Box plots depicting Piezo1 mCherry (left) and actin-GFP (right) front-back polarity in untreated (NT) and chemokine-stimulated conditions. **C.** Polar plots depicting relative Piezo1 mCherry and actin-GFP spatial distribution in Jurkat cells with respect to direction of cell trajectory in untreated (top panel) and SDF1α (bottom panel) treated cells. Spatial distribution was calculated for all the time points of time-lapse imaging (see methods).

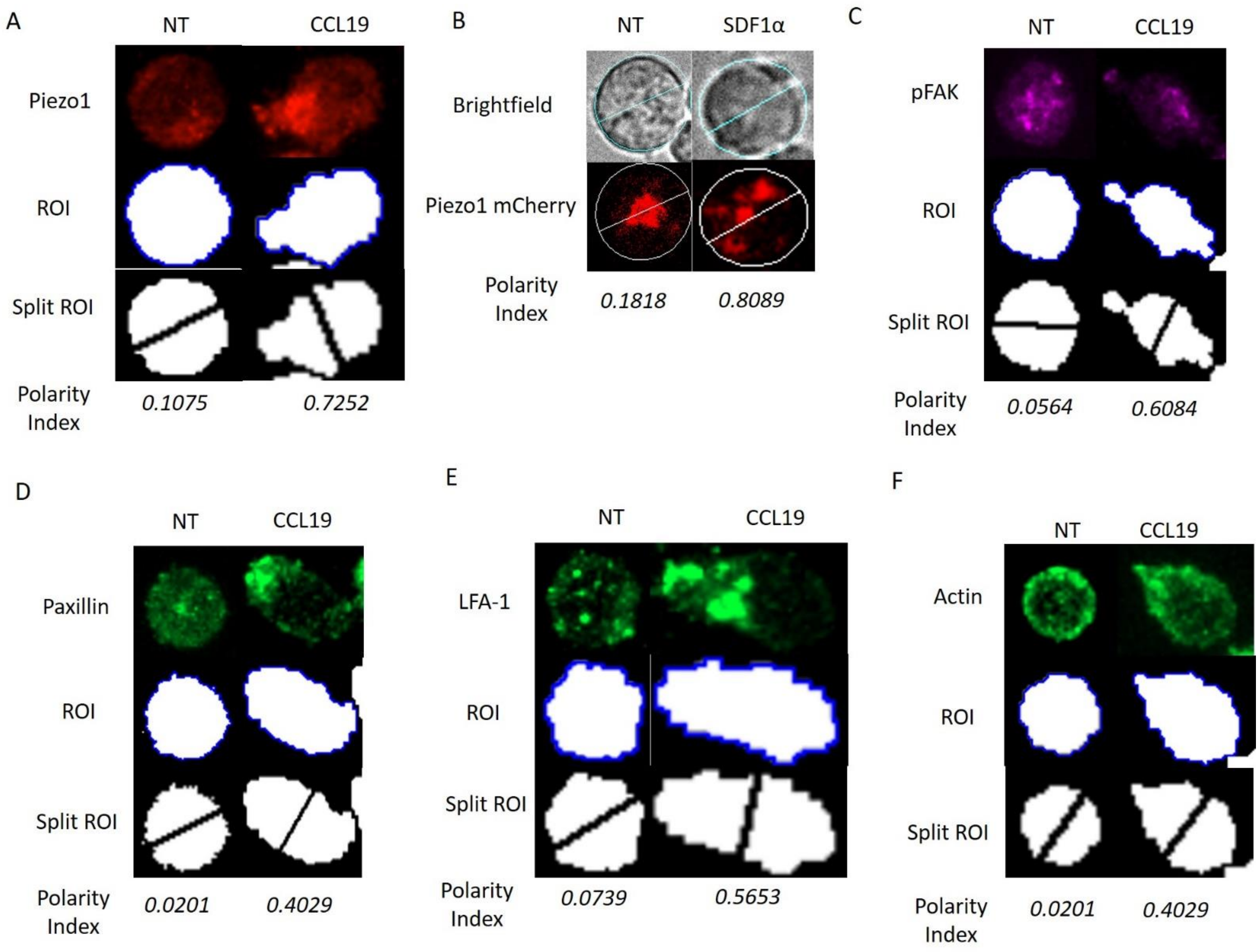


**Methods-figure supplement 1.** Representative analyses for calculation of cell polarity of **A.** Piezo1 (fixed); **B.** Piezo1 mCherry (live); **C.** phosphorylated FAK (pFAK, fixed cell); **D.** paxillin (fixed cell); **E.** CD11a/LFA1 (fixed cell); **F.** actin (fixed cell). Region-of-interest identifying cell boundary and polarity indices mentioned. All analyses performed on Fiji (see methods).

#### VIDEOS LEGENDS

**Video 1.** Representative video of cell trace-stained $CD4^+$ T lymphocytes transfected with GFP construct and Piezo1 siRNA, in the presence of recombinant human CCL19 on ICAM1-coated plates, for a total duration 15 minutes at 30 seconds per frame. The left panel represents superimposed cell trace and GFP signals of moving cells. The right panel shows tracks of the moving cells. Boxed cells are $GFP^+$ cells. 20X Magnification. Related to Figure 1A-C and Figure 1-figure supplement 1D.

**Video 2.** Representative 3D projection of Z-stacks of $CD4^+$ T lymphocyte treated with recombinant human CCL19 for 15 minutes, immunostained with Piezo1 antibody and Hoechst. 63X magnification. Related to Figure 1F.

**Video 3.** Representative immunofluorescence time-lapse video of Piezo1-mCherry expressing Jurkat cell moving on ICAM1-coated dish in the presence of recombinant human SDF1α. Images were acquired manually for a total duration of 15 minutes at 1 minute per frame interval. Images were captured at 40X magnification. Related to Figure 1I.

**Video 4.** Representative 3D rendering of confocal Z-stacks of $CD4^+$ T lymphocyte treated with recombinant human CCL19 for 10 minutes, immunostained with phosphorylated FAK antibody and Hoechst. 63X magnification. Related to Figure 3A.

**Video 5.** Representative 3D rendering of confocal Z-stacks of $CD4^+$ T lymphocyte depicting Piezo1 and CD11a/LFA-1 distribution, upon 30 minutes of recombinant CCL19 treatment. 63X magnification. Related to Figure 4A.

**Video 6.** Representative 3D rendering of confocal Z-stacks of control siRNA-transfected $CD4^+$ T lymphocyte depicting CD11a/LFA-1 polarity, upon 30 minutes of recombinant CCL19 treatment. 63X magnification. Related to Figure 4E.

**Video 7.** Representative 3D rendering of confocal Z-stacks of Piezo1 siRNA-transfected $CD4^+$ T lymphocyte depicting CD11a/LFA-1 polarity, upon 30 minutes of recombinant CCL19 treatment. 63X magnification. Related to Figure 4E.

**Video 8.** Representative 3D rendering of confocal Z-stacks of control siRNA-transfected $CD4^+$ T lymphocyte depicting phosphorylated Akt polarity, upon 30 minutes of recombinant CCL19 treatment. 63X magnification. Related to Figure 4F.

**Video 9.** Representative 3D rendering of confocal Z-stacks of Piezo1 siRNA-transfected $CD4^+$ T lymphocyte depicting phosphorylated Akt polarity, upon 30 minutes of recombinant CCL19 treatment. 63X magnification. Related to Figure 4F.

**Video 10.** Representative 3D rendering of confocal Z-stacks of $CD4^+$ T lymphocyte depicting Piezo1 and actin distribution, upon 30 minutes of recombinant CCL19 treatment. 63X magnification. Related to Figure 6A.

**Video 11.** Representative 3D rendering of confocal Z-stacks of control siRNA-transfected $CD4^+$ T lymphocyte depicting actin polarity, upon 30 minutes of recombinant CCL19 treatment. 63X magnification. Related to Fig. 6B.

**Video 12.** Representative 3D rendering of confocal Z-stacks of Piezo1 siRNA-transfected $CD4^+$ T lymphocyte depicting loss of actin polarity, after 30 minutes of recombinant CCL19 treatment. 63X magnification. Related to Fig. 6B.

**Video 13.** Representative time-lapse video of Piezo1-mCherry/actin-GFP expressing Jurkat cell, moving on ICAM1-coated dish in the presence of recombinant human SDF1α. Images were acquired as a Z-stack (1µm per stack) at 63X/1.40 oil magnification. Total duration of time-lapse was 5 minutes at 30 seconds per frame. Images are represented as maximum intensity projection of the Z stack. Related to Figure 6F.

#### SUPPLEMENTARY FILES

**Supplementary File 1.** Supplementary File 1 includes a detailed list of key resources used in the study.

**Supplementary File 2:** Supplementary File 2 is an excel file containing all data corresponding to each figure mentioned in the study.

**Supplementary File 3:** Supplementary File 3 includes a zip file containing codes used in Fiji (.ijm files) and MATLAB (.m files) image analysis.
