## Supplementary figures and images for "Piezo1 mechanosensing regulates integrin-dependent chemotactic migration in human T cells"

### Supplemental video 1

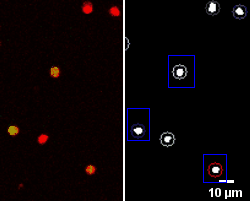

### Supplemental video 2

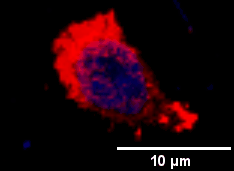

### Supplemental video 3

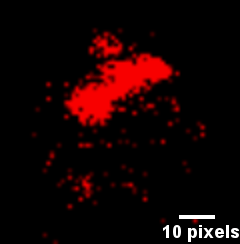

### Supplemental video 4

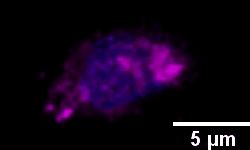

### Supplemental video 5

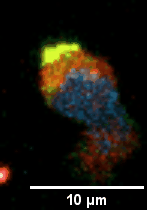

### Supplemental video 6

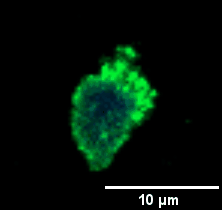

### Supplemental video 7

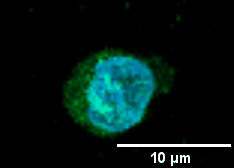

### Supplemental video 8

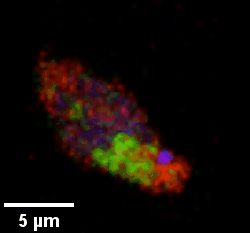

### Supplemental video 9

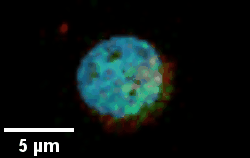

### Supplemental video 10

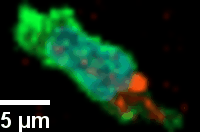

### Supplemental video 11

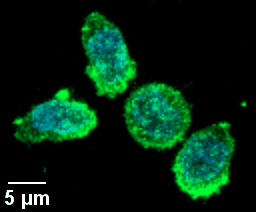

### Supplemental video 12

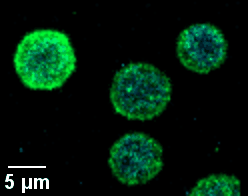

### Supplemental video 13

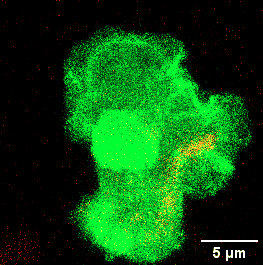
